# Transmission-blocking activity of artesunate, chloroquine and methylene blue on *Plasmodium vivax* gametocytes

**DOI:** 10.1101/2024.02.18.580875

**Authors:** Victor Chaumeau, Praphan Wasisakun, James A. Watson, Thidar Oo, Sarang Aryalamloed, Mu Phang Sue, Gay Nay Htoo, Naw Moo Tha, Laypaw Archusuksan, Sunisa Sawasdichai, Gornpan Gornsawun, Somya Mehra, Nicholas J. White, François H. Nosten

## Abstract

*Plasmodium vivax* is now the main cause of malaria outside Africa. The gametocytocidal effects of antimalarial drugs are important to reduce malaria transmissibility, particularly in low transmission settings, but they are not well characterized for *P. vivax*. The transmission-blocking effects of chloroquine, artesunate and methylene blue on *P. vivax* gametocytes were assessed. Blood specimens were collected from patients presenting with vivax malaria, incubated with or without the tested drugs, and then fed to mosquitos from a laboratory-adapted colony of *Anopheles dirus* (a major malaria vector in Southeast Asia). The effects on oocyst and sporozoite development were analyzed under a multi-level Bayesian model accounting for assay variability and the heterogeneity of mosquito *Plasmodium*-infection. Artesunate and methylene blue, but not chloroquine, exhibited potent transmission-blocking effects. Gametocyte exposures to artesunate and methylene blue reduced the mean oocyst count 469 fold (95%CI: 345 to 650) and 1438 fold (95%CI: 970 to 2064) respectively. The corresponding estimates for the sporozoite stage were a 148 fold reduction (95%CI: 61 to 470) and a 536 fold reduction (95%CI: 246 to 1311) in the mean count, respectively. In contrast, high chloroquine exposures reduced the mean oocyst count by only 1.40 fold (95%CI: 1.20 to 1.64) and the mean sporozoite count 1.34 fold (95%CI: 1.12 to 1.66). This suggests that patients with vivax malaria often remain infectious to anopheline mosquitos after treatment with chloroquine. Immediate initiation of primaquine radical cure or use of artemisinin combination therapies would reduce the transmissibility of *P. vivax* infections.

## INTRODUCTION

*Plasmodium vivax* is a major cause of malaria worldwide. Approximately one third of the global population is at risk of infection. There are about10 million symptomatic cases each year (1). Vivax malaria has been relatively neglected because it rarely causes acute death (2,3), although it is associated with indirect morbidity, poor pregnancy outcomes and, in highly endemic areas, recurrent infections contribute to anaemia-related mortality (4,5). *Plasmodium vivax* is associated with repeated relapses from persistent liver stages (hypnozoites) and is particularly difficult to control and eliminate (6,7). Treatment of symptomatic malaria with effective antimalarials reduces transmission. This plays a central role in malaria control and elimination in the low transmission settings where *P. vivax* is prevalent (8). In *Plasmodium falciparum* infections, gametocytogenesis is delayed, so prompt effective treatment reduces transmissibility (9). The treatment of vivax malaria is more complex. Schizonticidal drugs (artemisinin-based combination therapies or chloroquine) active against the pathogenic asexual blood stages are used to clear parasitaemia and obtain clinical remission, but to achieve radical cure (i.e. killing the hypnozoites and thereby preventing subsequent relapses) treatment with an 8-aminoquinoline (primaquine or tafenoquine) is required in addition (10). In contrast to *P. falciparum,* gametocytogenesis in *P. vivax* infections occurs together with asexual stage development, so symptomatic patients are usually infectious to vector anopheline mosquitos.

The inability to cryopreserve and then conduct long-term culture of *P. vivax* compromises laboratory assessment of transmission-blocking activity outside endemic areas. These assessments therefore require the proximity of insectary, laboratory and parasitaemic patients. As a result, few studies have been performed, and the effects of antimalarial drugs on *P. vivax* gametocytes are not well characterised (11). Chloroquine is considered active against *Plasmodium* gametocytes except for the mature stages of *P. falciparum* (12). However, transmission of *P. vivax* to mosquitos has been observed for up to 72 hours after starting treatment which suggests that chloroquine may lack activity against mature *P. vivax* gametocytes (13). With the exception of the 8-aminoquinolines, artemisinins are more active against mature *Plasmodium* gametocytes than other antimalarials (14). In vivax malaria the artemisinin combination therapy (ACT) dihydroartemisinin-piperaquine was reported to have a superior transmission-blocking effect compared with chloroquine, but the individual effects of the two drugs in the combination were not studied (15). Similarly, the artesunate-mefloquine ACT regimen co-administered with primaquine was reported recently to have a superior transmission-blocking effect compared with chloroquine co-administered with primaquine or tafenoquine (16). This supports earlier observations that artemisinin derivatives had greater activity than chloroquine in reducing gametocyte carriage in vivax malaria (17,18). High concentrations of methylene blue, which has a potent gametocytocidal activity in *P. falciparum* (19,20), were shown recently to block transmission of *P. vivax* gametocytes in membrane feeding experiments but the sample size was very small (only five patients were recruited in this study) (21).

The aim of this study was to compare the transmission-blocking activity of artesunate, chloroquine and methylene blue on *P. vivax* gametocytes. *Anopheles dirus* mosquitos (a major vector in Southeast Asia) from a laboratory-adapted colony were fed on blood specimens collected from vivax malaria patients and incubated with or without drug. Drug effects on oocyst and sporozoite counts in mosquito samples were analyzed under a Bayesian multi-level negative-binomial model accounting for assay variability and heterogeneity of *Plasmodium*-development in the mosquito (22).

## RESULTS

Overall 38 adult vivax malaria patients provided blood samples, 342 *Anopheles dirus* mosquito batches were fed on these samples, and 20,908 mosquitos were dissected for assessment of either oocyst or sporozoite counts (Appendix, Table S1). Baseline sample characteristics (i.e., on the day of sample collection, before the 24-hour incubation with or without drug) are shown in Table 1. Overall, the median asexual parasite and gametocyte densities were 13,161 parasites/µL (inter-quartile range, IQR: 6981 to 27,798) and 1092 gametocytes/µL (IQR: 473 to 2009) respectively. All but one of the blood samples were infectious to mosquitos (the sample that was not infectious at baseline became infectious after 24 hours of incubation). The median oocyst index (i.e., the proportion of mosquitos harbouring malaria oocysts per batch) was 0.96 (IQR 0.84 to 0.98) and the median oocyst count in mosquitos was 63.8 oocysts per mosquito (IQR 4.1 to 124.9). The corresponding figures for the sporozoite stage were 0.7 (IQR 0.35 to 0.9) and 211 sporozoites per mosquito (IQR 5 to 4321). The median ratio of the median parasite count in the controls after 24 hours of incubation without drug to the median baseline parasite count (on the day of sample collection) was 0.93 (IQR: 0.51 to 3.68) and 0.9 (IQR: 0.01 to 6.56) for the oocyst and sporozoite stages, respectively (Appendix, Figure S1).

**Table 1.** Characteristics of the blood samples at baseline.

| Characteristic | Value in the experimental group |  |
| --- | --- | --- |
|  | Median | IQR |
| No. of blood samples |  |  |
| Artesunate | 9 | - |
| Chloroquine | 21 | - |
| Methylene blue | 8 | - |
| Asexual parasitaemia (no. asexual parasites /μL of blood) | | |
| Artesunate | 12,109 | 8258 to 16,608 |
| Chloroquine | 12,740 | 3610 to 21,652 |
| Methylene blue | 17,718 | 10,116 to 28,675 |
| Gametocytemia (no. asexual parasites /μL of blood) | | |
| Artesunate | 767 | 524 to 1350 |
| Chloroquine | 898 | 195 to 1849 |
| Methylene blue | 1821 | 1309 to 4063 |
| Oocyst index |  |  |
| Artesunate | 0.98 | 0.94 to 0.98 |
| Chloroquine | 0.96 | 0.68 to 0.98 |
| Methylene blue | 0.96 | 0.92 to 0.98 |
| Median oocyst count (no. oocysts per mosquito) |  |  |
| Artesunate | 95 | 48.5 to 127 |
| Chloroquine | 18.5 | 2 to 92 |
| Methylene blue | 114.5 | 54.6 to 183.9 |
| Sporozoite index |  |  |
| Artesunate | 0.60 | 0.10 to 0.70 |
| Chloroquine | 0.65 | 0.27 to 0.86 |
| Methylene blue | 0.95 | 0.87 to 1.00 |
| Median sporozoite count (no. sporozoites per mosquito) |  |  |
| Artesunate | 33 | 0 to 211 |
| Chloroquine | 122 | 2.5 to 2377 |
| Methylene blue | 4604 | 2402 to 10,876 |

411 Abbreviations: IQR, inter-quartile range.

Chloroquine exhibited little transmission-blocking activity on *P. vivax* gametocytes, despite the high concentrations used (Figure 1). Of all dissected mosquitos in the chloroquine spiked samples, 2974/4036 (74%) carried oocysts in the treated replicates compared with 3299/4026 (82%) in the controls (relative risk, RR: 0.85 [95% confidence interval, CI: 0.82 to 0.88], p<0.0001) and 701/1177 (60%) carried sporozoites in the treated replicates compared with 785/1228 (64%) in the controls (RR: 0.89 [95%CI: 0.81 to 0.97], p=0.005). In contrast, artesunate and methylene blue almost completely interrupted gametocyte transmission. For artesunate, only 207/1798 (12%) of the mosquitos carried oocysts in the treated replicates compared with 1591/1797 (89%) in the controls (RR: 0.057 [95%CI: 0.044 to 0.075], p<0.0001) and 1/360 (0.3%) dissected mosquitos carried sporozoites in the treated replicates versus 152/360 (42%) in the controls (RR: 0.005 [95%CI: 0.001 to 0.035], p<0.0001); for methylene blue, only 76/1599 (5%) carried oocysts in the treated replicates versus 1470/1592 (92%) in the controls (RR: 0.039 [95%CI: 0.028 to 0.054], p<0.0001) and only 5/320 (1.6%) carried sporozoites in the treated replicates versus 267/320 (83%) in the controls (RR: 0.01 [95%CI: 0.003 to 0.027], p<0.0001). The lower sporozoite index observed in the controls of the artesunate group in comparison to the sporozoite index at baseline was probably explained by the detrimental effect of artesunate wash off on sporogony (see model coefficient estimate below).

**Figure 1.**
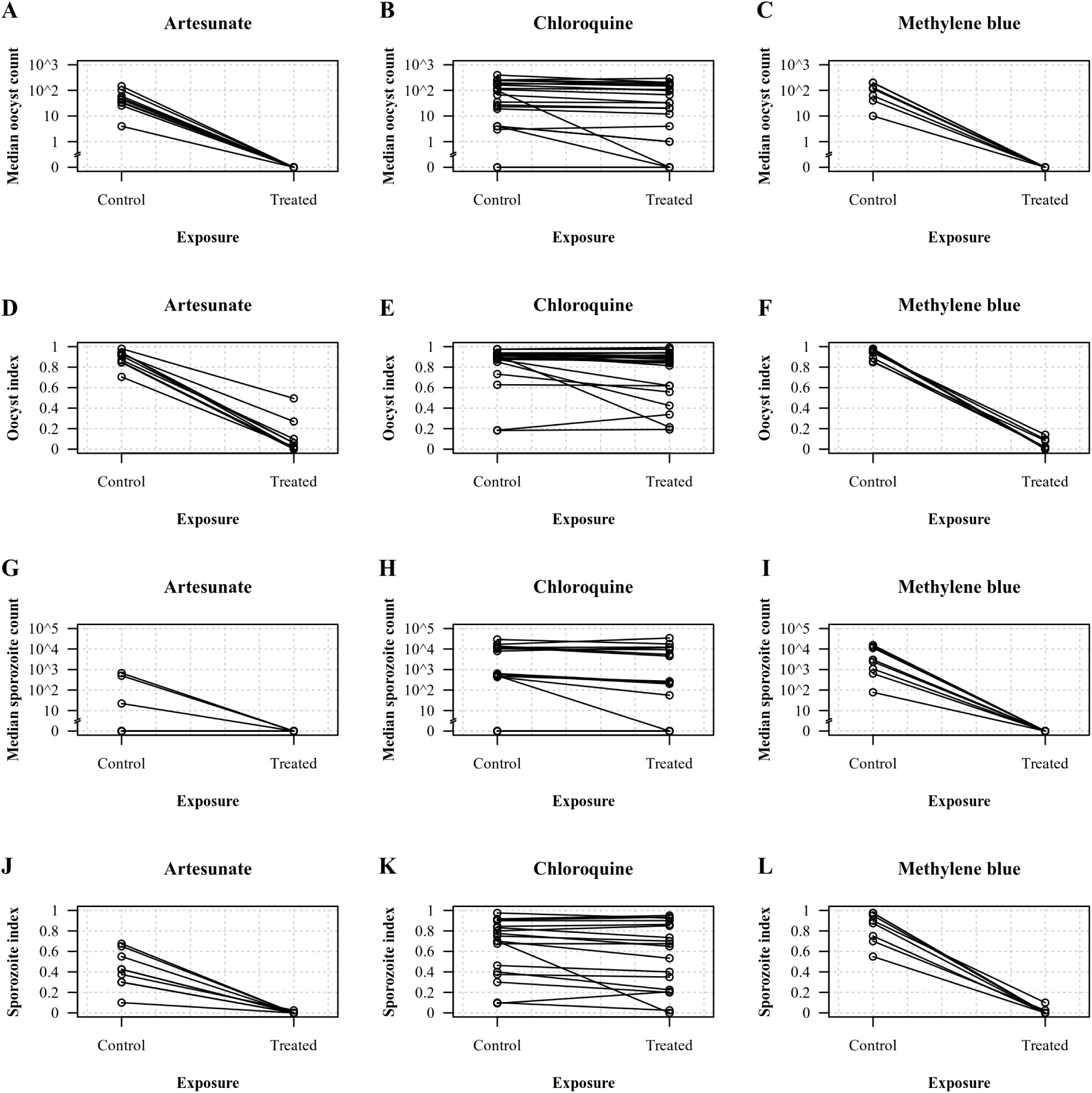
Effects of chloroquine, methylene blue and artesunate on the development of *P. vivax* in *An. dirus*. (A-C) Median oocyst count; (D-F) oocyst index; (G-I) median sporozoite count; (J-L) sporozoite index. Values in the control and treated replicates were collated by assay run.

As observed previously, there was considerable heterogeneity in the count data across mosquitos within a batch, and considerable variability in the count data across blood samples and experimental batches. To account for this heterogeneity, we estimated the drug effects under a Bayesian multi-level model (mixed effects) whereby the count data were modelled as negative binomial with the dispersion parameter as a parametric function of the mean count (Table 2; see Methods) (22). In contrast to the previous data describing proportions of mosquitos with parasites, the model parameterized the drug effect as a reduction in the mean number of parasites per mosquito, accounting for variability across blood samples and mosquito batches. Under this model, gametocyte exposure to chloroquine decreased the mean oocyst count by only 1.40 fold (95% credible interval, CrI: 1.20 to 1.65; observed mean of 100 oocysts per mosquito in the controls versus 69 in the treated replicates), and it decreased the mean sporozoite count by 1.34 fold (95%CrI: 1.12 to 1.66; observed mean of 14,414 sporozoites per mosquito in the controls versus 11,132 in the treated replicates). In contrast, artesunate reduced the mean oocyst count by 469 fold (95%CrI: 345 to 650; observed mean of 60 oocysts per mosquito in the controls versus 0.22 in the treated replicates), and methylene blue reduced the oocyst count 1438 fold (95%CrI: 970 to 2064; observed mean of 107 oocysts per mosquito in the controls versus 0.08 in the treated replicates). For the sporozoite counts the estimates were a 148-fold reduction for artesunate (95%CrI: 61 to 470; observed mean of 1303 sporozoites per mosquito in the controls versus 0.1 in the treated replicates), and a 536 fold for methylene blue (95%CrI: 246 to 1311; observed mean of 13,914 sporozoites per mosquito in the controls versus 1.8 in the treated replicates). The model fitted the data well as shown by the inferred relationship between the mean parasite count and proportion of *Plasmodium*-infected specimens in mosquito samples (Figure 2). As expected, both inter- and intra-experiment variability were large and inter-experiment variability was larger than intra-experiment variability. The median fold-variation in the mean parasite count across blood samples was 1.07 (IQR: 0.62 to 2.08, range: 0.16 to 4.00) and 1.32 (IQR: 0.47 to 2.40, range: 0.12 to 6.83) fold for the population means for the oocyst and sporozoite stages, respectively (Appendix, Figure S2). The median fold-variation in the mean parasite count across technical replicates was 1.00 (IQR: 0.77 to 1.30, range: 0.005 to 4.05) and 0.99 (IQR: 0.94 to 1.05, range: 0.62 to 1.88) fold the for the patient mean for the oocyst and sporozoite stages, respectively (Appendix, Figure S3). One sample with abnormally high intra-experiment variability in the mean oocyst count was detected but no obvious explanation for this outlier was identified. Inclusion or exclusion of this sample from the analysis did not significantly change the results (data not shown). Moreover, the development of sporozoites mirrored that of the oocysts: a 10-fold increase in the mean oocyst count was associated with a 3.52-fold (95%CI: 2.15 to 4.90-fold) increase in the mean sporozoite count (Appendix, Figure S4). To explain variation in blood meal infectiousness to mosquitos across blood samples, the log_10_[mean oocyst count], log_10_[asexual parasitaemia] and log_10_[gametocytaemia] assessed on admission (i.e., on the collection day before the 24-hour incubation time with or without drug) were introduced as linear predictors of the mean parasite count in mosquito samples of the experimental replicates (i.e., after 24 hours of incubation with or without drug). A 10-fold increase in the mean oocyst count and gametocytaemia at baseline were associated with a 1.76 [95%CrI: 1.20 to 2.48] and a 6.47-fold increase [95%CrI: 3.04 to 13.37] respectively in the mean oocyst count in the experimental replicates; there was no significant association between the mean oocyst count in experimental replicate and baseline asexual parasitaemia (model coefficient estimate: 1.08 [95%CrI: 0.56 to 2.11]) or artesunate wash off (model coefficient estimate: 0.84 [95%CrI: 0.41 to 1.71]). A 10-fold increase in baseline gametocytaemia and artesunate wash off were respectively associated with a 3.56-fold increase [95%CrI: 1.21 to 9.20] and a 3.85-fold decrease (model coefficient estimate: 0.26 [95%CrI: 0.10 to 0.61]) in the mean sporozoite count in the experimental replicates. There was no significant association between the mean sporozoite count in the experimental replicates and the mean oocyst count (model coefficient estimate: 1.21 [95%CrI: 0.72 to 1.96]) or asexual parasitaemia (model coefficient estimate: 1.22 [95%CrI: 0.48 to 2.96]) at baseline.

**Figure 2.**
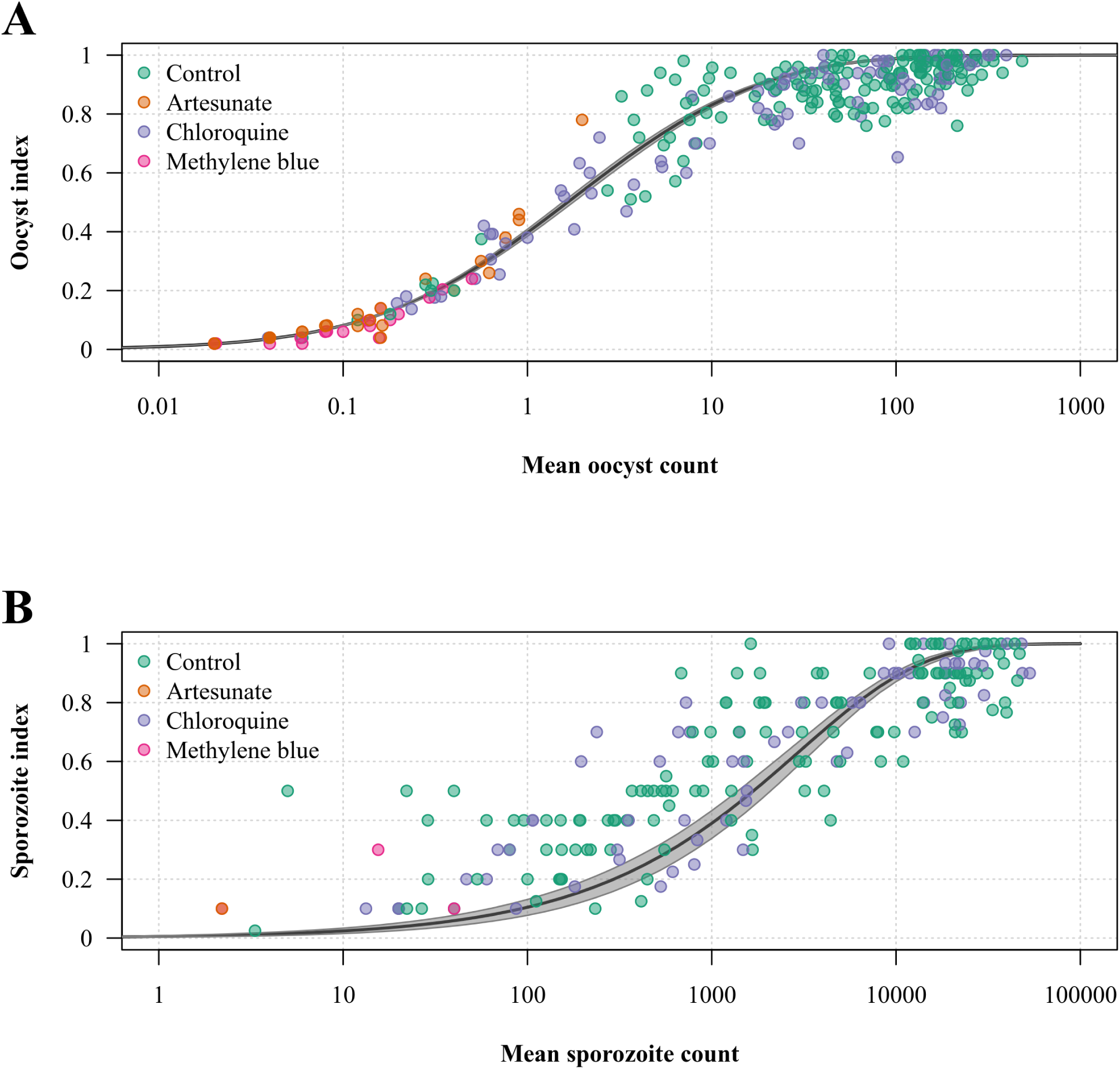
Relationship between the mean parasite count and proportion of *Plasmodium*-infected mosquitos in the assay. (A) Paired mean number of oocysts per mosquito and oocyst index determined in the control and treated replicates; (B) paired mean number of sporozoites per mosquito and sporozoite index. The black line and shaded area show the model-fitted relationship plotted using a_0_ and a_1_ estimates given by the model output and the corresponding 95% credible interval, respectively.

**Table 2.** Parameter estimates given by the output of transmission-blocking activity model.

| Parameter | Oocysts |  | Sporozoites |  |
| --- | --- | --- | --- | --- |
|  | Median of posterior draws | 95%CrI | Median of posterior draws | 95%CrI |
| $\mu_{\text{population}}$ | 44.6 | 31.3 to 63.7 | 4906.6 | 2875 to 8158.8 |
| $\sigma_{\text{patient}}$ | 0.92 | 0.74 to 1.15 | 1.22 | 0.98 to 1.54 |
| $\sigma_{\text{batch}}$ | 0.46 | 0.41 to 0.53 | 0.31 | 0.23 to 0.41 |
| $a_0$ | 0.41 | 0.37 to 0.45 | 0.0007 | 0.0004 to 0.0012 |
| $a_1$ | 0.19 | 0.16 to 0.21 | 0.61 | 0.56 to 0.68 |
| Artesunate effect | 0.0021 | 0.0015 to 0.0029 | 0.0068 | 0.0021 to 0.0165 |
| Chloroquine effect | 0.71 | 0.61 to 0.84 | 0.74 | 0.60 to 0.90 |
| Methylene blue effect | 0.0007 | 0.0005 to 0.0010 | 0.0019 | 0.0008 to 0.0041 |
| $\log_{10}$[Baseline oocyst count] | 1.76 | 1.20 to 2.48 | 1.21 | 0.72 to 1.96 |
| $\log_{10}$[Baseline parasitemia] | 1.08 | 0.56 to 2.11 | 1.22 | 0.48 to 2.96 |
| $\log_{10}$[Baseline gametocytemia] | 6.47 | 3.04 to 13.37 | 3.56 | 1.21 to 9.20 |
| Wash off | 0.84 | 0.41 to 1.71 | 0.26 | 0.10 to 0.61 |

Abbreviation: CrI, credible interval.

## DISCUSSION

This assessment of the transmission-blocking effects of antimalarial drugs on *P. vivax* gametocytes revealed that chloroquine has little activity against *P. vivax* gametocytes. Its activity is probably limited to the immature forms having a food vacuole, i.e., the pre-macrogametocytes originally described by Boyd (23), whereas high doses of artesunate and methylene blue have potent *P. vivax* gametocytocidal and thus transmission-blocking effects.

The data also confirm previous observations on the relationship between intensity (number of oocysts or sporozoites per mosquito) and prevalence (oocyst or sporozoite index) of *Plasmodium*-infections in artificially infected mosquitos (22). Transmission-blocking drugs primarily reduce the intensity (number and viability) of oocyst development and the resulting effect on prevalence varies with the mean parasite count in mosquito samples, being less in high-intensity than in low-intensity infections. This is well described by a negative binomial model with a dispersion parameter as a function of the mean count (22). As oocysts arising from gametocytes exposed to an antimalarial drug may fail to produce viable sporozoites, the primary outcome of transmission-blocking assays should therefore be the reduction in sporozoite carriage.

The results highlight differences in the intrinsic susceptibility of *P. vivax* and *P. falciparum* gametocytes to antimalarial drugs. Unlike other human malaria parasite species, *Plasmodium falciparum* gametocytes’ emergence is delayed with respect to asexual parasite densities, their maturation takes longer, and mature stage V gametocytes are intrinsically resistant to most antimalarial drugs, except methylene blue and the 8-aminoquinolines (24). Artesunate, which kills young circulating sexual stages but fails to kill mature *P. falciparum* gametocytes (14), exhibited a potent transmission-blocking effect on *P. vivax*.

This study had several limitations. The characteristics of *P. vivax* gametocyte maturation and the determinants of gametocyte infectiousness to mosquitos are not well characterized (25). The sex- and stage-specific effects of drugs on the gametocytes were not assessed. Sex and stage-specific gametocytocidal effects were previously reported with *P. falciparum* (14,26). To the best of our knowledge, this has never been assessed in *P. vivax* probably because these aspects of *Plasmodium* biology are less well known in *P. vivax* than in P*. falciparum*. Drug concentrations were high, and the concentration-response relationships were not evaluated. The experiment was designed to maximize the power to detect a drug effect, and so a single high concentration was investigated. The correlation between the exposures of drugs *in vitro* and *in vivo* is also not well characterized and the concentrations tested in this study may not represent drug activity at therapeutic doses. Testing of lower concentrations of the active drugs would be informative. 8-aminoquinolines are considered potent gametocytocides, but the absence of ex-vivo metabolism precluded investigation of these prodrugs. Several approaches have been proposed to investigate drug metabolites *in vitro* including direct synthesis of stable metabolites identified during pharmacokinetic studies *in vivo* or *in situ* metabolism of the parent compound in the assay (27,28). Interestingly, primaquine, which has potent effects against *P. falciparum* gametocytes (29,30), was shown in one study to be less effective in killing *P. vivax* gametocytes (31). Assessment of the gametocytocidal effects of biotransformed primaquine and other 8-aminoquinolines on *P. vivax* gametocytes will require further research. Sporozoite viability was not assessed in this study and may lead to underestimation of the effect of chloroquine. However, successful invasion of the mosquito salivary glands is already an indication of their viability. This limitation could be addressed by assessing the development of liver stages inoculated with sporozoites detected in the assay (32). Susceptibility of asexual parasites to the drugs was not determined. It could be argued that the observed low transmission-blocking activity of chloroquine against *P. vivax* gametocytes results from parasite resistance rather than intrinsic lack of gametocyte susceptibility to the drug. However, this is unlikely given the good treatment efficacy and the reported data on *P. vivax* asexual blood stages susceptibility to antimalarial drugs in this study area (33).

Using gametocytocidal drugs (artemisinin combination treatments) for the first line treatment of vivax malaria may reduce infection transmissibility but it is important to consider the timing of gametocyte development and transmission *in vivo*. *Plasmodium vivax* gametocytes can arise directly from exo-erythrocytic schizonts and appear in the peripheral circulation as early as the asexual blood stages (32,34). In addition, the lower limit of gametocyte density for transmission to mosquito is lower in *P. vivax* than other human malaria parasite species: successful transmission to vector mosquitos can occur with densities of gametocytes as low as 5 gametocytes per μL (35). These densities are below the limit of routine microscopy detection. Previous exposure increases the pyrogenic threshold (circa 10 parasites per μL in naive individual versus approximately 200 parasites per μL in the immune) (36,37) and infected individuals can bear transmissible densities of gametocytes without any symptom. Therefore, in endemic areas, the majority of patients are infectious to mosquitos before diagnosis and treatment of the infection (17,38). Nevertheless, if artemisinin-based combination therapies are indeed superior to chloroquine in preventing *P. vivax* transmission as this study suggests, this is an additional argument in favour of a unified treatment for all malarias (39), particularly if radical treatment is delayed or not given.

## MATERIAL AND METHODS

### Participants and sample collection

Patients with vivax malaria attending outpatient consultation at the clinics of the Shoklo Malaria Research Unit in Wang Pha and Maw Ker Tai (Northwest border of Thailand) were invited to participate in the study by giving a single 10-mL blood sample drawn into a sterile sodium heparin tube before receiving antimalarial drug treatment. The sample was kept into a Thermos® bottle filled with water warmed at 37°C until processing (typically within 1 hour after collection). The study was approved by the Oxford Tropical Research Ethics Committee, the Tak Public Health Office Ethics Committee and the Tak Province Border Community Ethics Advisory Board (40). All participants provided their written informed consent to participate in the study.

In order to estimate parasite densities on admission, a thin smear and a thick film of participant blood sample were prepared on a glass slide, stained with 5% Giemsa for 35 min and examined under a microscope at a 1000 magnification using standard procedures (41), and a complete blood count was performed. The proportion of red blood cells infected with malaria parasites was estimated by recording the total parasite count in 2000 red cells in the thin smear. If no parasite was detected in 2000 red cells (3/38 samples), the count was determined for 500 white cells in the thick film. Then, gametocyte and asexual parasites were counted separately in a subset of 100 parasites and the proportions were used to estimate gametocytaemia and asexual parasitaemia from the total parasite count per 2000 red cells or per 500 white cells and the concentration of red cells or white cells in participant blood sample, as appropriate. All slides were read independently by two blinded microscopists and discrepant results were resolved by a third microscopist. The mean values of the two concordant readings were used in the analysis.

### Compounds

Chloroquine and artesunate were supplied by the Worldwide Antimalarial Resistance Network (WWARN). Chloroquine diphosphate (Sigma-Aldrich, catalog no. C6628) stock solution was prepared at a concentration of 97 mmol/L in water. Artesunate (Sigma-Aldrich, catalog no. 88495-63-0) stock solution was prepared at a concentration of 52 mmol/L in 100% ethanol. Methylene blue (Poveblue®, methylthioninium chloride trihydrate solution at 5 mg/mL or 13 mmol/L) was kindly provided by Provepharm (Marseille, France) and used as a stock solution. All stock solutions were kept at -80°C, used within 6 months and thawed only once before being used in the assay.

### Parasite culture

The blood sample was transferred into a 50-mL conical tube and centrifuged at 500 g for 5 minutes at 37°C. The serum and buffy coat were discarded, and the cell pellet was washed twice with 45 mL of incomplete culture medium warmed at 37°C using the same centrifugation conditions. Incomplete culture medium was composed of RPMI-1640 (Sigma-Aldrich, catalog no. R6504) supplemented with 2 g/L of NaHCO_3_ (Sigma-Aldrich, catalog no. S6014), 5.7 g/L of HEPES (Sigma-Aldrich, catalog no. H4034) and 18 mg/L of hypoxanthine (Sigma-Aldrich, catalog no. H9636). The cell pellet was resuspended into complete culture medium warmed at 37°C to in a total volume of 20 mL. The complete culture medium was composed of incomplete medium supplemented with 10% of heat-inactivated AB serum. The serum was inactivated by heating at 56°C for 30 min, aliquots were kept at -80°C and thawed only once before performing the assay. Eight culture flasks containing 8 mL of complete culture medium were warmed at 37°C without (control flasks, n = 4) or with a spike of the test drug (treated flasks, n = 4) and were inoculated with 2 mL of the blood cell suspension (total volume of 10 mL). The flasks were incubated in with 5% CO_2_ at 37°C for 24 hrs. An additional wash off step was added for assay runs carried out with artesunate to mimic the rapid elimination of this drug *in vivo*. After 4 hours, the contents of all flasks (both control and treated states) were transferred into 15-mL conical tubes and washed twice with 12 mL of complete culture medium using the same centrifugation conditions, then resuspended into 10 mL of complete culture medium, and then incubated for a further 20 hrs.

### Assay design, sample size and power

Mosquitos from a laboratory-adapted colony of *An. dirus* were artificially infected with *P. vivax* by carrying out membrane feeding experiments using the vivax malaria blood samples. The mosquito colony was maintained as described previously (42). Before the test feed, the blood specimen was incubated with artesunate (1 μmol/L for 4 hrs, followed by 20 hrs of incubation without drug), chloroquine (5 μmol/L for 24 hrs) or methylene blue (1 μmol/L for 24 hrs). The same specimen incubated without drug was used as the control. In assay runs carried out with artesunate, all control and treated flasks were washed to control for the effects of washing steps on sample infectiousness to mosquitos. Four technical replicates were performed for each group (treated and control), yielding 8 mosquito batches per assay run. The artesunate and methylene blue concentrations each of 1 μmol/L were chosen to represent the high concentrations typically used for *in vitro* drug screening; chloroquine was tested at a concentration of 5 μmol/L because a concentration of 1 μmol/L did not exhibit evident transmission-blocking activity during preliminary experiments in the initial assay setup (data not shown). Drugs were assigned to blood samples in sequential order: the assay was repeated 18 times with chloroquine, 8 times with methylene blue and 9 times with artesunate. The assay was then repeated 3 times with a different batch of chloroquine to exclude assessment bias relating to compound quality. The development of oocysts was assessed in samples of 50 mosquitos per batch 7 days after the feed, yielding a total of 450 oocyst counts per experiment: 50 for the baseline feed, 200 in the controls and 200 in the treated replicates. Similarly, the development of sporozoites was assessed in samples of 5 mosquitos per batch 14 and 15 days after the feed (10 mosquitos per batch in total), yielding a total of 90 sporozoite counts per experiment: 10 for the baseline feed, 40 in the controls and 40 in the treated replicates. The sporozoite count could not be determined in three assay runs because the laboratory shut down during a COVID-19 outbreak. To estimate the required sample size, a multi-level Bayesian model was fitted to a data set of oocyst counts in 97 artificial mosquito infections carried out at the same facility and the model output was used to perform simulation experiments. Power to detect a 10% reduction in the mean oocyst count was calculated at varying numbers of dissected mosquitos per technical replicate, number of technical replicates per assay run and number of independent assay runs. Using this power calculation, the study was powered to detect a 10% reduction in the mean oocyst count with 7 independent assay runs for each drug.

### Membrane-feeding assay

At the end of incubation, the content of the flasks was transferred into 15-mL conical tubes and centrifuged at 500 g for 5 min at 37°C. The supernatant was discarded and the cell pellet (approximately 500 μL) was resuspended into 500 μL of heat-inactivated AB serum warmed at 37°C. The suspension was then fed to the *An. dirus* mosquitos with a Hemotek membrane feeding system (Blackburn, United Kingdom) using 1-mL reservoirs covered with stretched Parafilm (Bemis, USA). The assay was carried out with 5-7 day-old nulliparous female imagoes starved by removing the wet towel covering the cage and the sugar source for 4 to 6 hrs before the feed. Mosquitos were transferred into 750 cm^3^ plastic containers at a density of 150 specimens per cup and left undisturbed for 30 minutes before the feed; eight cups were prepared in total (one per replicate) and the same mosquito batch was used for a given assay run. The feed was carried out by putting the Hemotek insert on top of the corresponding mosquito container and regularly blowing through the net every five minutes for 1 hour. Fully engorged mosquitos were transferred into 4500 cm^3^ plastic containers at 15-minute intervals until 100 fully engorged mosquitos per replicate were collected (typically about 1 hour). Engorged mosquitos were kept at 25°C and provided with 10% sugar solution *ad libitum* until dissection.

### Assessment of oocyst and sporozoite development

Dissected mosquito midguts were stained with 2% Mercurochrome solution for 5 min, observed under a microscope at a 40 magnification and the number of oocysts per midgut was recorded. Pairs of salivary glands were crushed in 1 μL of 1X PBS using the corner of a glass slide. The crushed salivary glands were rinsed with 20 μL of 1X PBS. The mixture (approximately 15 uL) was then transferred into 1.5 mL plastic tubes and kept on wet ice until determination of the sporozoite concentration with a hemocytometer (typically within 4 hrs after the dissection). If no sporozoite was detected in the hemocytometer, the dried slide was examined under a microscope at a 40 magnification to identify mosquito specimens that carried few sporozoites, below the detection limit of the hemocytometer. The sporozoite count in such specimens was arbitrarily set to 10 sporozoites per mosquito.

### Data analysis

The proportion of *Plasmodium*-infected mosquitos was analyzed under a multi-level logistic regression model including group allocation as a linear predictor and a random effect across participant blood samples to account for correlation in mosquito *Plasmodium*-infection between experimental replicates of the same sample. The relative risk was then calculated using odds ratio estimate and proportion of infected specimens in the controls. Parasite count data (the number of oocysts and the sporozoites per mosquito) were analyzed under a Bayesian multi-level model taking into account intra- and inter-experiment variability as per Medley *et al.* (22). In order to consider heterogeneity of *Plasmodium* development in the mosquito, the likelihood function was a Negative Binomial distribution parameterized by its mean μ and the dispersion κ for integer parasite counts y; y ∼ Negative Binomial(μ, κ), with κ set as a function of the mean:

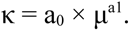

Under the Negative Binomial model, the prevalence of infection P (oocyst or sporozoite index, defined as the number of *Plasmodium*-infected specimens divided by the number of dissected specimens) varies as a function of the mean infection intensity:

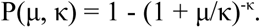

For each patient i and technical replicate k, the model predicted mean log parasite count μ_i,k_ was expressed as the sum of a patient dependent random effect ʎ_i_ and a batch random effect ʎ_k_; ʎ_i_ ∼ Normal(μ_i_, σ_patient_) and ʎ_k_ ∼ Student-t(7, 0, σ_batch_). The Student-t distribution with 7 degrees of freedom was chosen to accommodate the observed heterogeneity across batches (43). Thus, μ_i,k_ = ʎ_i_ + ʎ_k_ where μ_i_ is the mean log parasite count in mosquito samples fed on blood from patient i, such as μ_i_ ∼ Normal(μ_population_, σ_patient_), with μ_population_ being the mean log parasite count in mosquito samples fed on blood specimens from the overall patient population, σ_patient_ the standard deviation of individual patient mean log counts around the population mean and σ_batch_ the standard deviation of batch effects.

For a given blood sample, the drug treatment effect in treated replicates β_T[i]_ was parameterized in the model as a proportional decrease in the mean number of counts on the log scale. Thus, the likelihood of the count data y_i,k,T[i]_ (patient i, technical replicate k, treatment assignment T[i]) given the parameters is:

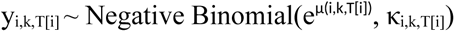

where μ_i,k,T[i]_ = ʎ_i_ + ʎ_k_ + β_T[i]_ + β_cov[i]_; and where κ_i,k,T[i]_ is a function of μ_i,k,T[i]_ as above. The additional model coefficients β_cov[i]_ accounts for the differences in experimental conditions for the artesunate samples (washing) and baseline characteristics of the sample (asexual parasitaemia, gametocytaemia and oocyst count assessed on the day of sample collection, before incubation with or without the test drug). Continuous covariables were log-transformed with a logarithm of base 10, meaning that a 10-fold increase in the covariable of interest was associated with a fold variation in the parasite count equal to the exponent of the coefficient estimate.

We used weakly informative priors to help computational convergence. These were μ_population_ ∼ Normal(5, 5) and μ_population_ ∼ Normal(9, 5) in the model fitted to oocyst and sporozoite data, respectively. The priors for other parameters were the same in both fits: σ_patient_ ∼ zero-truncated Normal(1, 0.25), σ_batch_ ∼ zero-truncate Normal(0.5, 0.25), log(a_0_) ∼ Normal(-1, 1), log(a_1_)∼ Normal(-1, 1), β_T[i]_ ∼ Normal(0, 1) and β_condition[i]_ ∼ Normal(0, 1). The model was run with 4 independent chains each consisting of 4000 iterations. Convergence of the chains was assessed by examining the values of effective sample size and Rhat and the traceplots (Appendix, Figure S5-8).

## DATA AVAILABILITY STATEMENT

All analysis code and data are available via an accompanying github repository: https://github.com/victorSMRU/transmission-blocking-plasmodium-vivax.

## ACKNOWLEDGEMENTS

We are very grateful to the volunteers who participated in this study. We thank the staff of the Entomology, Laboratory, Medical and Data Management Departments of the Shoklo Malaria Research Unit for their help with collection, processing and management of the samples and data included in this study. We thank The WorldWide Antimalarial Resistance Network (WWARN) for providing antimalarial drugs. We thank Dr. Georges Snounou for his kind help with literature review. The Shoklo Malaria Research Unit is part of the Mahidol-Oxford Research Unit, supported by Wellcome, U.K. (#220211). This research was funded by Wellcome. A CC BY or equivalent licence is applied to the author accepted manuscript arising from this submission, in accordance with the grant’s open access conditions.

