## Appendix for "Transmission-blocking activity of artesunate, chloroquine and methylene blue on *Plasmodium vivax* gametocytes"

**Table S1. Summary of assay runs.**

| Drug | No. blood samples | No. mosquito batches | | | No. dissected guts | | | No. dissected pairs of salivary glands | | |
|---|---|---|---|---|---|---|---|---|---|---|
| | | Base. | Cont. | Treat. | Base. | Cont. | Treat. | Base. | Cont. | Treat. |
| Artesunate | 9 | 9 | 36 | 36 | 450 | 1797 | 1798 | 90 | 360 | 360 |
| Chloroquine | 21 | 21 | 84 | 84 | 1008 | 4026 | 4036 | 263 | 1228 | 1177 |
| Methylene blue | 8 | 8 | 32 | 32 | 404 | 1592 | 1599 | 80 | 320 | 320 |

Abbreviation: base., baseline; cont., control; treat., treated.

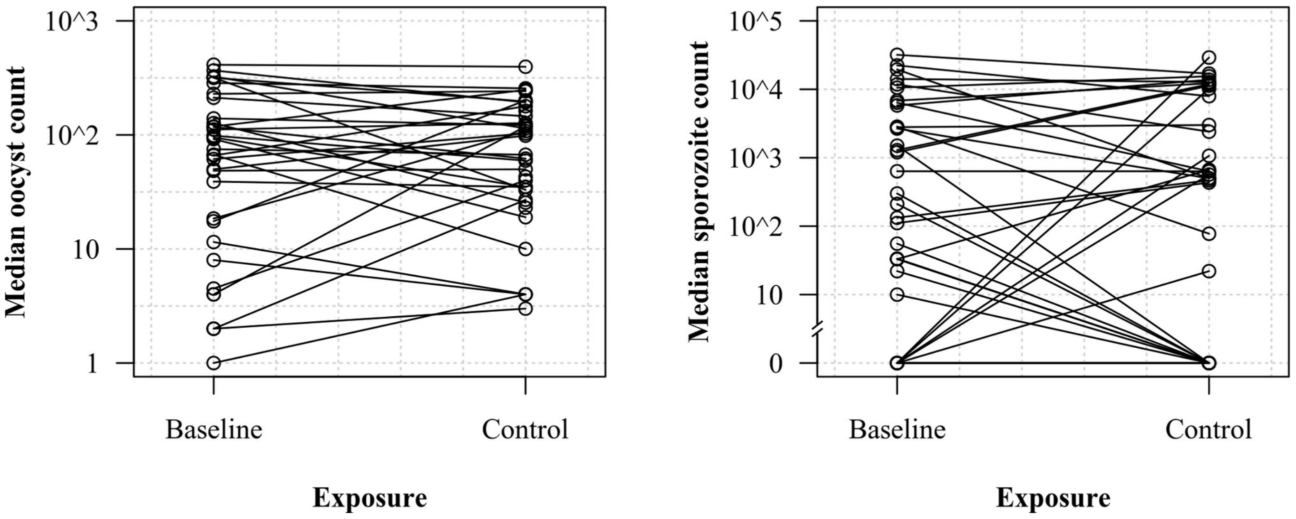


**Figure S1. Evolution of sample infectiousness to mosquitos during incubation *ex vivo*.** Variations in the median oocyst (left panel) and median sporozoite count (right panel) between baseline (i.e., on the day of sample collection, before incubation) and control (i.e., after 24 hours of incubation without drug) mosquito batches. Values of experimental replicates were collated by assay run.

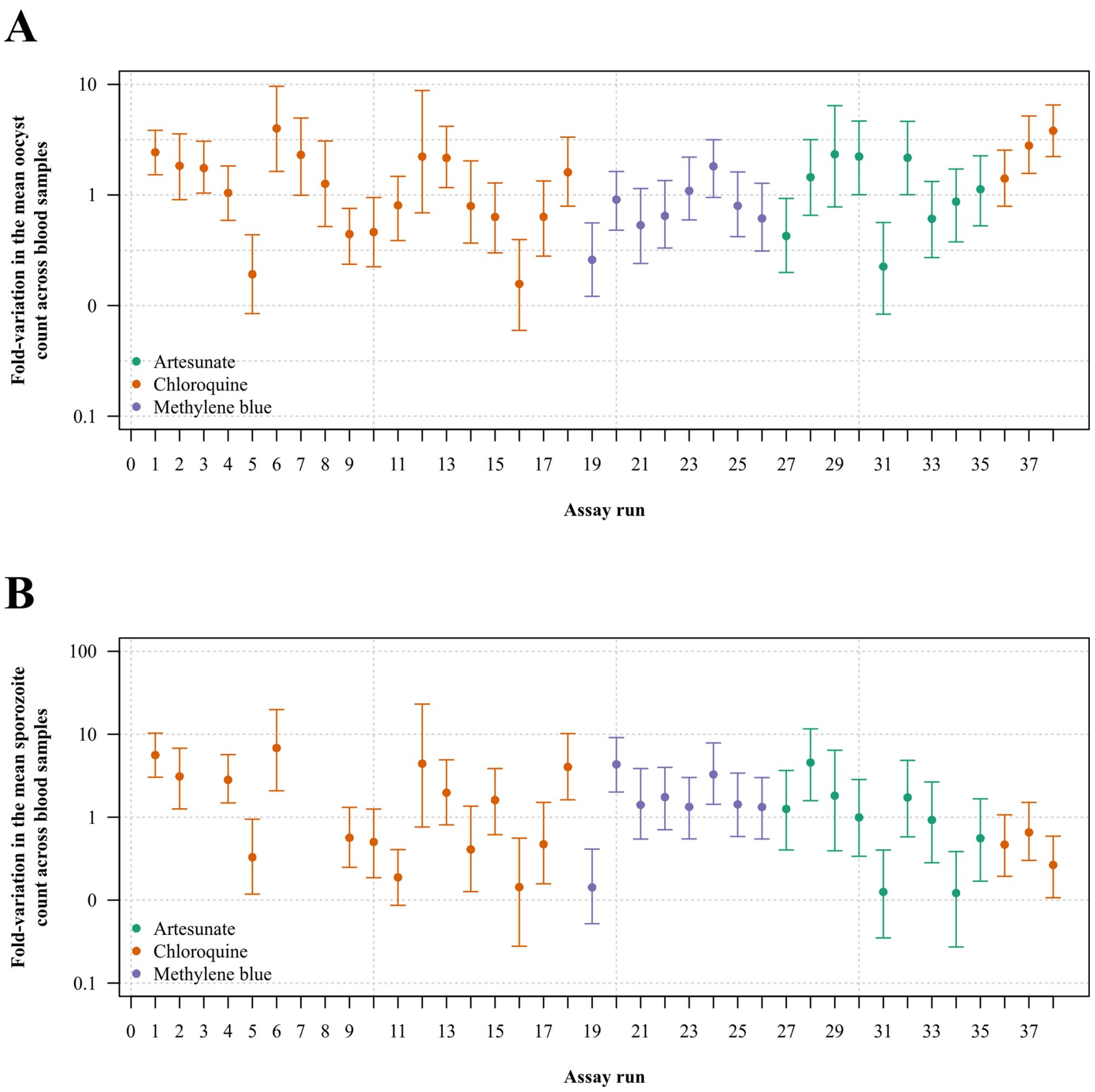


**Figure S2. Inter-experiment variability.** Mean estimated oocyst (top panel) and sporozoite (bottom panel) counts (in the untreated state) under the Bayesian multi-level model; points and error bars show the median and 95% credible interval of posterior draws, respectively.

A

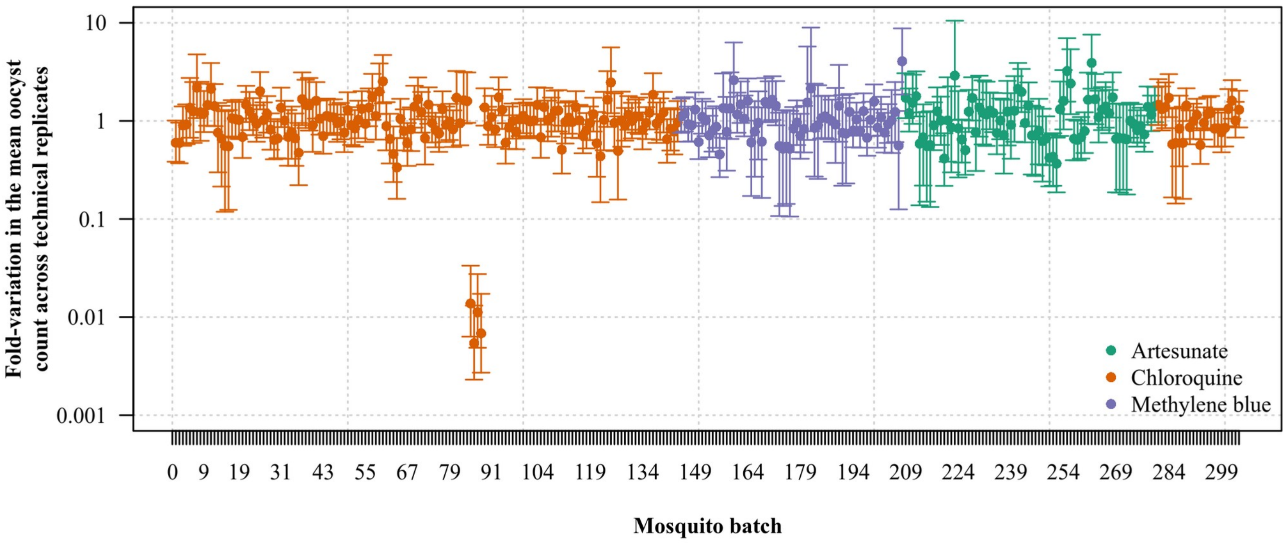


B

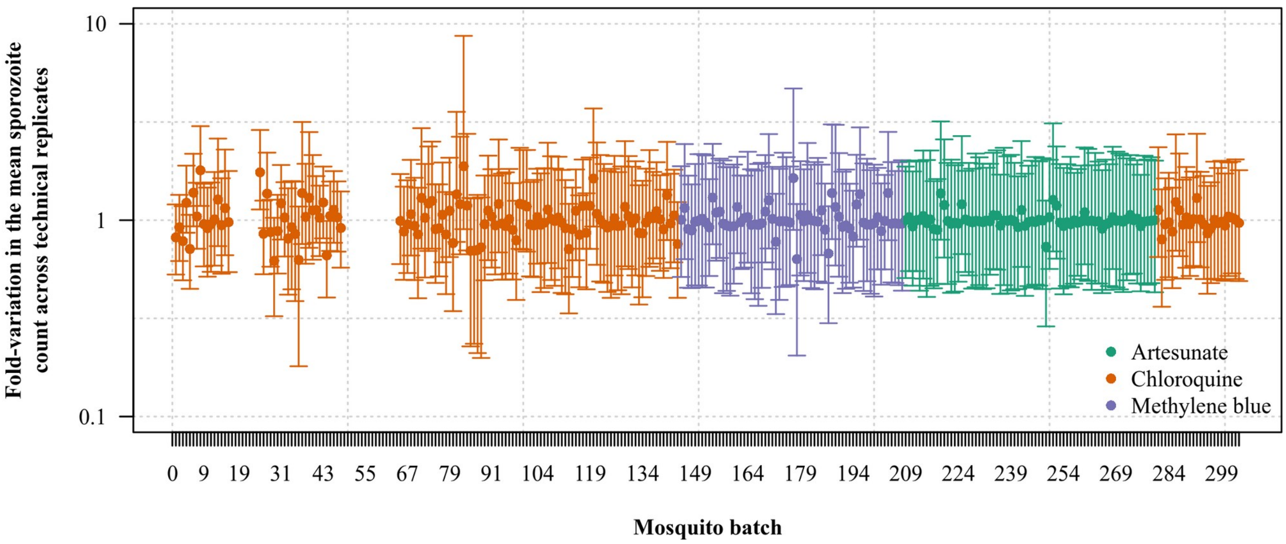


**Figure S3. Temporal trend in intra-assay variability.** Random variation of the mean oocyst (top panel) and sporozoite (bottom panel) count under the Bayesian multi-level model; points and error bars show the median and 95% credible interval of posterior draws, respectively.

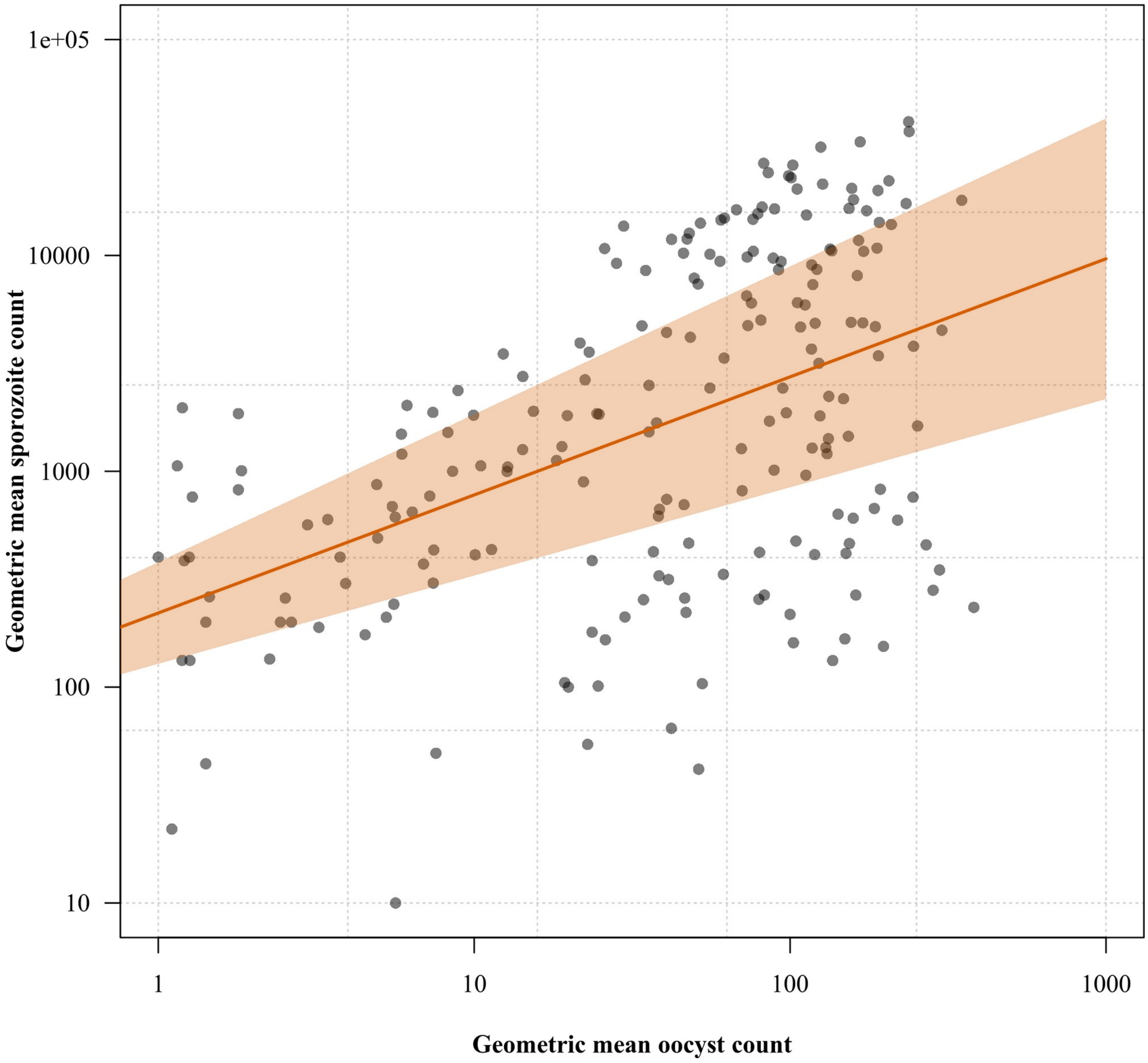


**Figure S4. Relationship between the mean oocyst and sporozoite count in the assay.** Points show the observed non-negative values plotted on the log scale; the orange line and shaded area show the linear regression fit and corresponding 95% confidence interval, respectively.

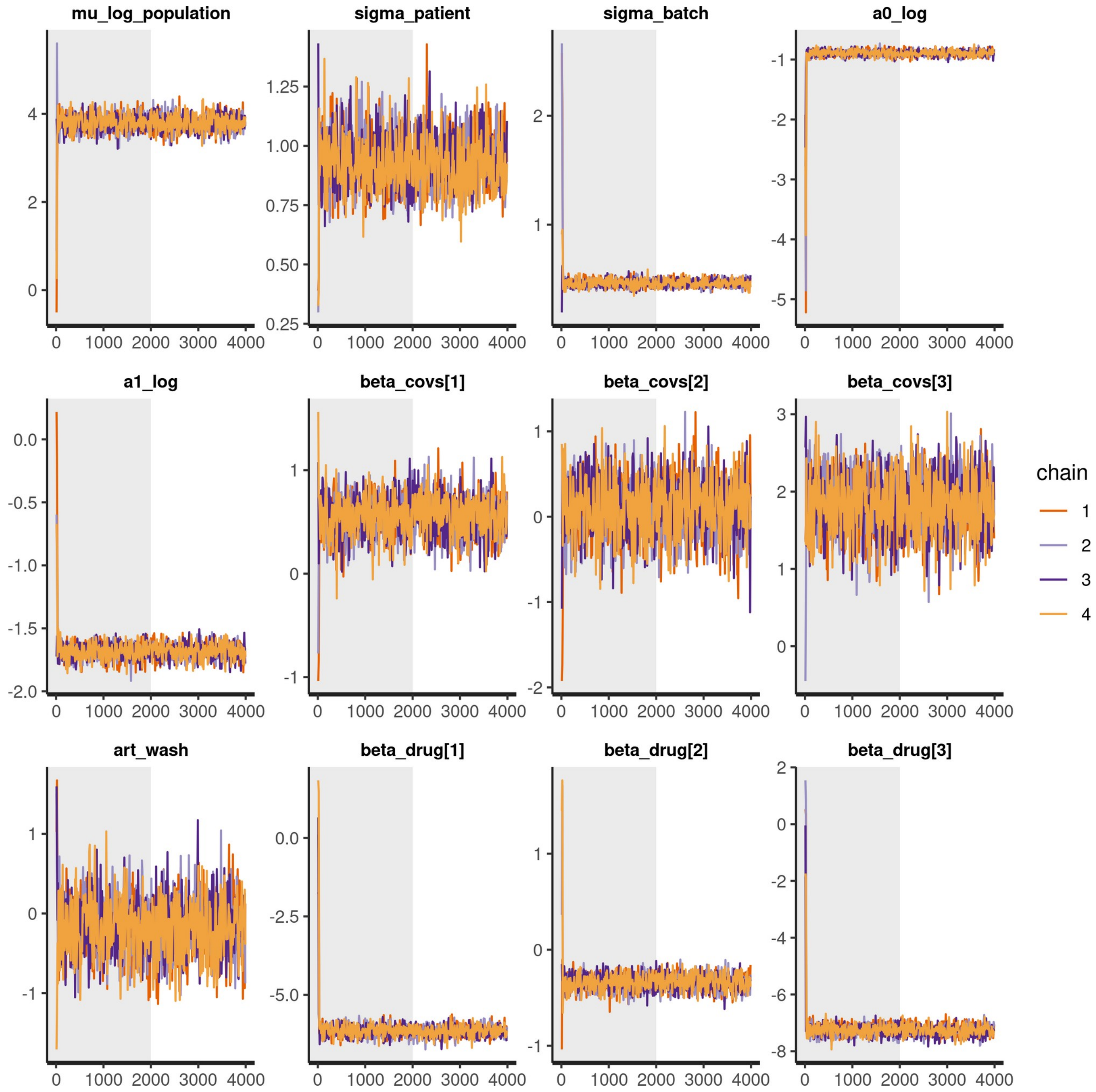


**Figure S5. Traceplot of MCMC chains showing mixing and convergence for the Bayesian multi-level model fitted to oocyst data.**

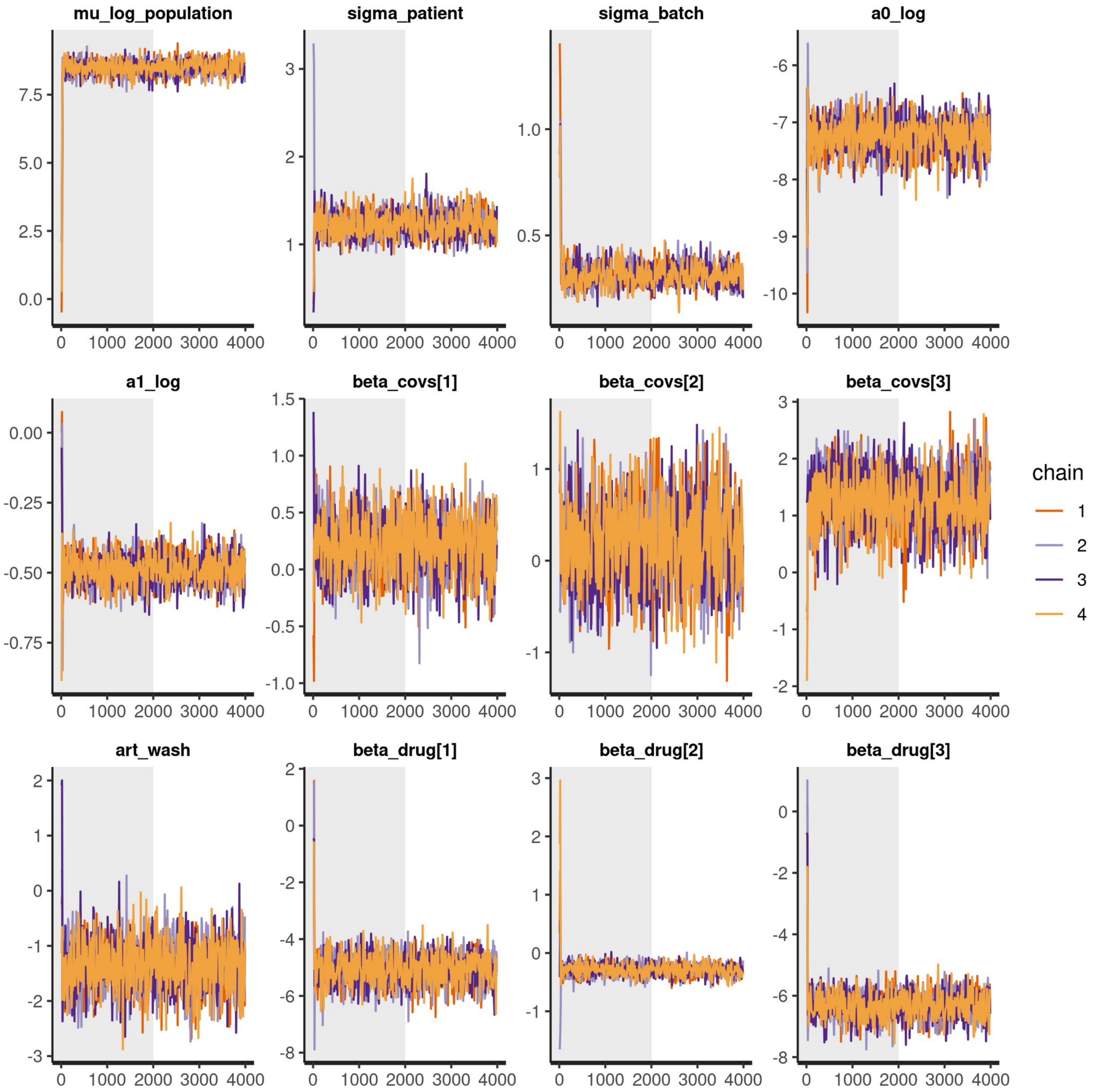


**Figure S6. Traceplot of MCMC chains showing mixing and convergence for the Bayesian multi-level model fitted to sporozoite data.**

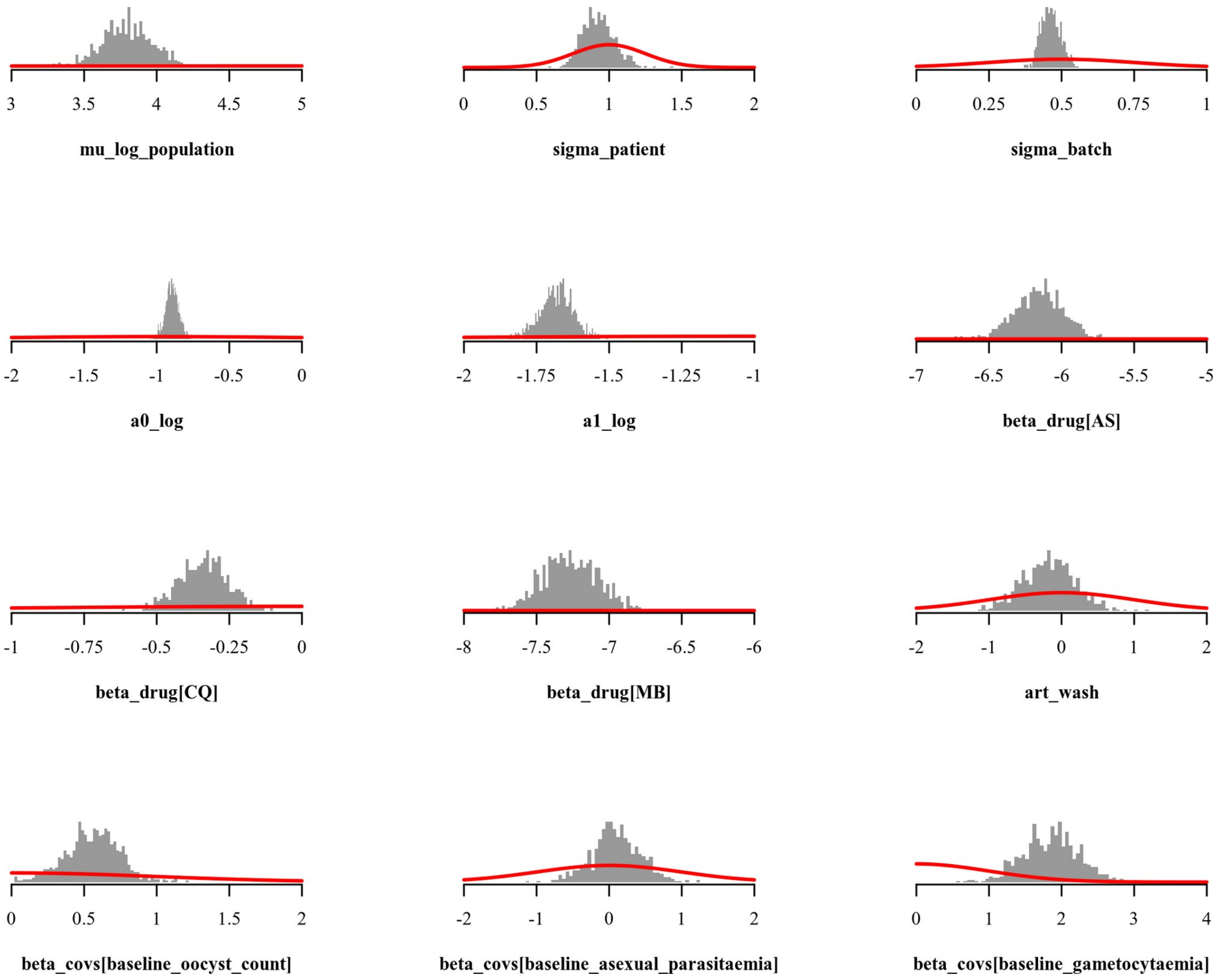


**Figure S7. Comparison between prior distributions (thick red lines) and posterior distributions (shown as histograms) for the parameters of Bayesian multi-level model fitted to oocyst data.**

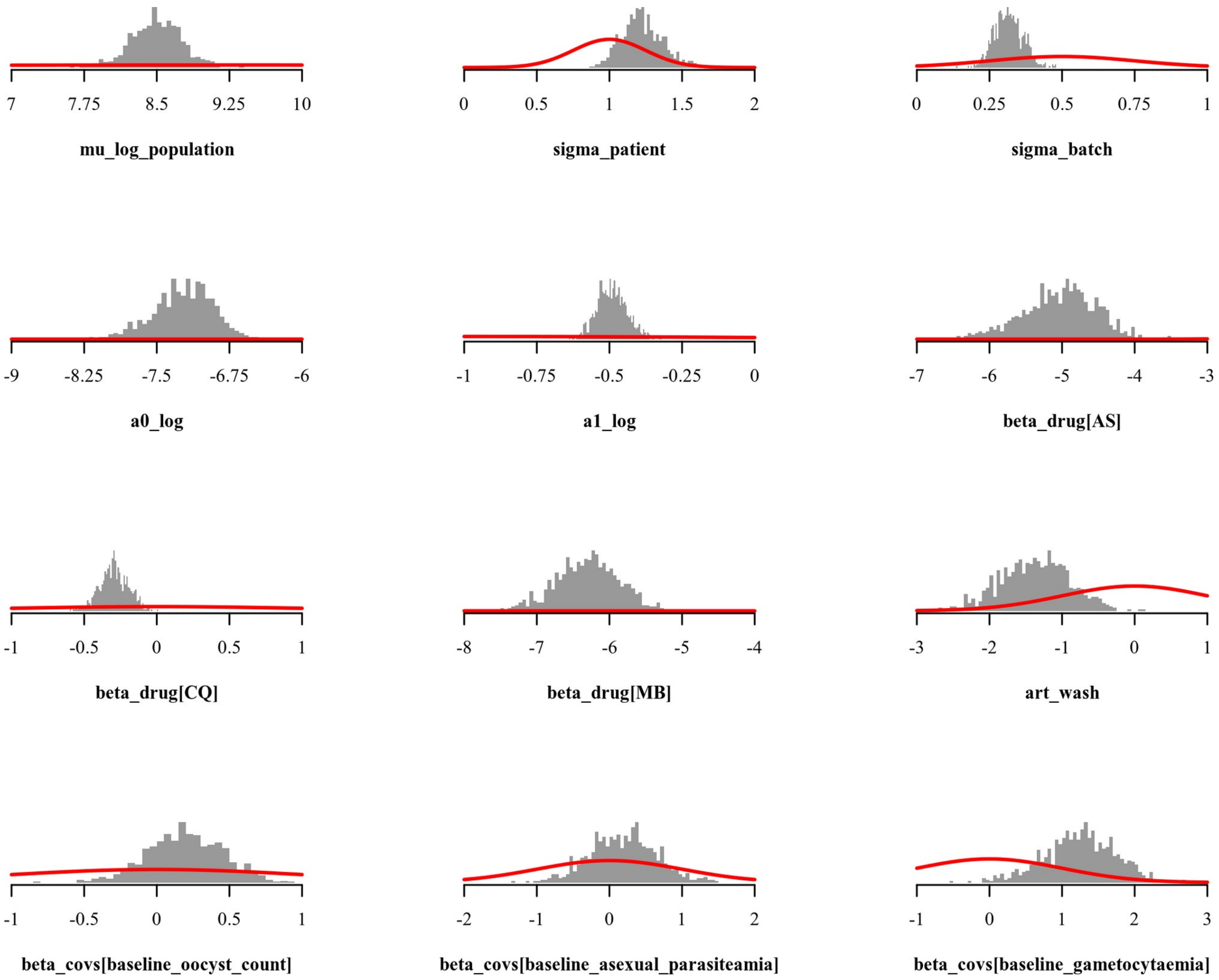


**Figure S8. Comparison between prior distributions (thick red lines) and posterior distributions (shown as histograms) for the parameters of Bayesian multi-level model fitted to sporozoite data.**
